# ClusterEnG: An interactive educational web resource for clustering big data

**DOI:** 10.1101/120915

**Authors:** Mohith Manjunath, Yi Zhang, Steve H. Yeo, Omar Sobh, Nathan Russell, Christian Followell, Colleen Bushell, Umberto Ravaioli, Jun S. Song

## Abstract

**Summary:** Clustering is one of the most common techniques used in data analysis to discover hidden structures by grouping together data points that are similar in some measure into clusters. Although there are many programs available for performing clustering, a single web resource that provides both state-of-the-art clustering methods and interactive visualizations is lacking. ClusterEnG (acronym for Clustering Engine for Genomics) provides an interface for clustering big data and interactive visualizations including 3D views, cluster selection and zoom features. ClusterEnG also aims at educating the user about the similarities and differences between various clustering algorithms and provides clustering tutorials that demonstrate potential pitfalls of each algorithm. The web resource will be particularly useful to scientists who are not conversant with computing but want to understand the structure of their data in an intuitive manner.

**Availability:** ClusterEnG is part of a bigger project called KnowEnG (Knowledge Engine for Genomics) and is available at http://education.knoweng.org/clustereng.

## 1 Introduction

Clustering is one of the most powerful and widely used analysis techniques for discovering structure in large data sets by grouping data points that are similar according to some measure. Several programming languages such as R (R Core Team, 2015) and Python (Pedregosa et al., 2011) offer libraries or packages for clustering custom data and generating static plots. However, interactive visualization, which aids the user in understanding the data at a deeper level, requires additional libraries and external software. Web servers, such as ClustVis (Metsalu et al., 2015), provide a simple yet powerful interface for visualizing Principal Component Analysis (PCA) and heatmap plots. However, at present, ClustVis limits the file upload size to 2 MB, and the plots are also static.

The advent of next-generation sequencing has enabled researchers to generate big data at an unprecedented rapid pace. Therefore, there is acute need for resources that can enable the users of high-dimensional biological data to quickly perform “first-hand” analysis, such as clustering (Stephens et al., 2015). The main challenges to building such a resource are handling big data and facilitating its interpretability. Client-side computer systems or web browsers may not always be powerful enough for efficient navigation through the data. The NIH has recently funded Big Data to Knowledge (BD2K) Centers to tackle this type of challenges. As part of the KnowEnG BD2K Center, we have developed a web-based resource called ClusterEnG (acronym for Clustering Engine for Genomics) for clustering big data with efficient parallel algorithms and software containerization. In order to facilitate the visualization of high-dimensional data, we implemented interactive versions of PCA, which is one of the most popular dimensional reduction techniques. ClusterEnG’s principal component plots in 2D and 3D allow intuitive exploration of structures in data. Fig. 1 illustrates the flowchart of various components of ClusterEnG, from user-uploaded data to output visualizations.

**Fig. 1.**
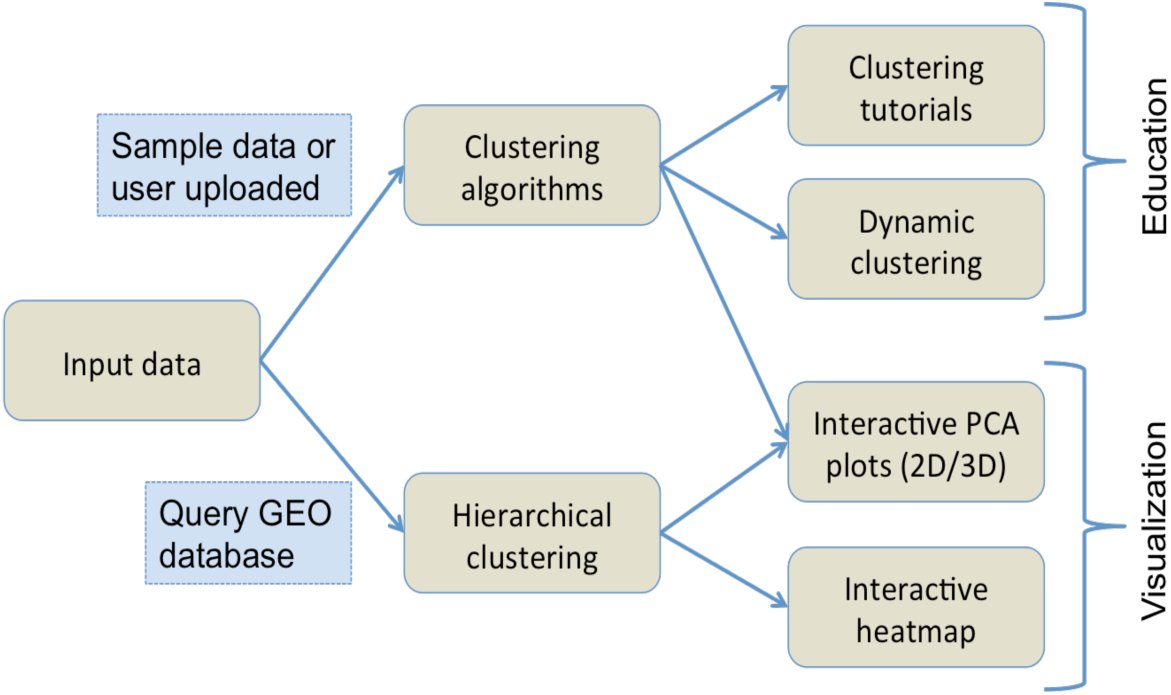
Typical workflow of ClusterEnG encompassing educational and visualization components.

## 2 Interface and Methods

### Input data and output

The user can upload custom data or choose one of the preloaded sample data sets for clustering. ClusterEnG accepts data in a tabular format of rows and columns, allowing the user to analyze most data sets generated by typical biological experiments, such as RNA-seq, microarray and drug-response data. The input data are then read in R utilizing the fast and convenient “fread” function from the *data.table* package (Dowle et al., 2014). The ClusterEnG server currently accepts files up to a size of 1GB, which will be steadily increased in the future. The uploaded file will be securely stored on the server for seven days, during which the user can retrieve the file or run more jobs from the same browser (with cookies enabled).

Currently, the server contains two public sample data sets: the gene expression data from NCI60 cancer cell lines (Ross et al., 2000) and B-cell lymphoma gene expression data (Alizadeh et al., 2000). The NCI60 data (9,707 genes, 64 samples) provide a good data set to explore and assess the quality of clustering from various algorithms implemented in ClusterEnG. The B-cell lymphoma data have a similar number of samples as the NCI60 data, but contains a larger number of genes (18,432 genes, 67 samples).

Clustering results are made available to the user in a CSV format in mainly two different ways. First, the user can download a single file with the entire data, sample/feature annotation and clustering results. Second, the user can select a subset of data interactively and download the clustering labels for the chosen data points. The user can also download snapshots of principal component plots in PDF, PNG and SVG formats.

### Clustering algorithms

ClusterEnG provides seven clustering algorithms, including parallel implementations for two algorithms. Currently, serial implementations are written in R programming language using various packages available in the CRAN repository (R Core Team, 2015). The seven algorithms include k-means, k-medoids, affinity propagation, spectral clustering, Gaussian mixture model, hierarchical clustering and DBSCAN (Ester et al., 1996). The parallel code for k-means algorithm utilizes a software package written in C (Liao, 2005), while parallel spectral clustering implements a C++ code (Chen et al., 2011). In addition to providing a choice of algorithms, the user is given a list of commonly used parameters for a subset of algorithms to modify and visualize the changes (Fig. 2).

**Fig. 2.**
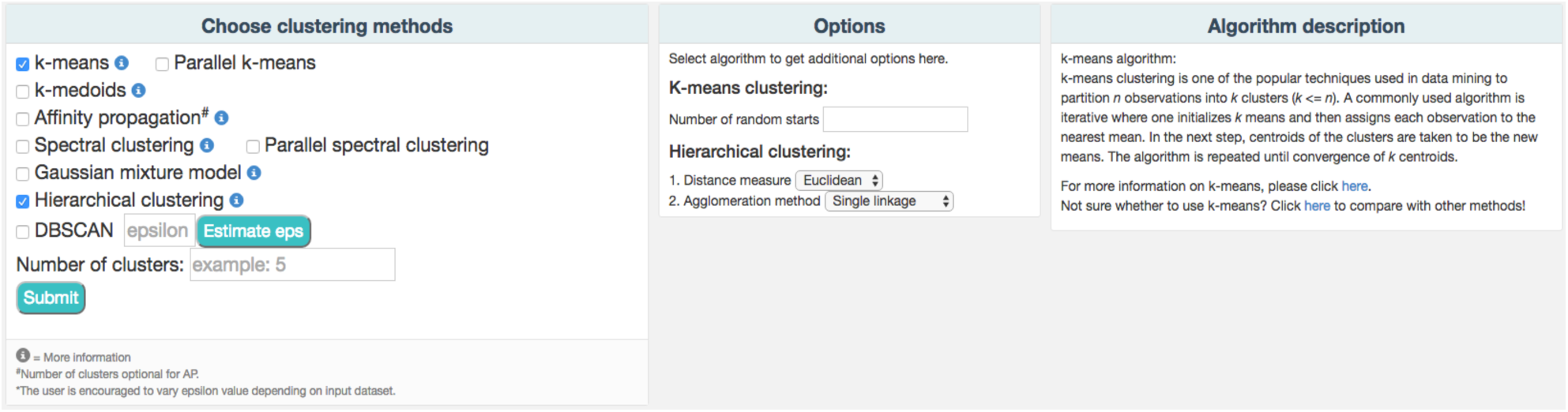
A partial snapshot of ClusterEnG user interface showing a choice of clustering algoritluns and related options.

ClusterEnG also features a module for querying the GEO database (Davis et al., 2007) to download data and draw an interactive heatmap with hierarchical clustering based on the InCHlib JavaScript library (Škuta et al., 2015).

### Docker containerization

We employ novel and efficient methods to handle the analysis of large files. The input data and user-selected algorithms from the front-end are dynamically packaged into a Docker container (Merkel, 2014) on the back-end wherein the code (serial or parallel) is executed and the results are returned to the main server. Chronos is used to schedule jobs by spawning Docker containers into an Apache Mesos cluster, which automatically utilizes available processors for parallel runs.

### Interactive 2D/3D visualization

After PCA is performed on data, projection coefficients on the first three principal components are used to generate three linked scatter plots for each pair of the components (Fig. 3a). Interactive plots are displayed using JavaScript library d3.js (Bostock et al., 2011) and jQuery to allow zooming, group selecting, mousing-over for annotation, and highlighting a region/cluster which maps to other PC direction plots.

**Fig. 3.**
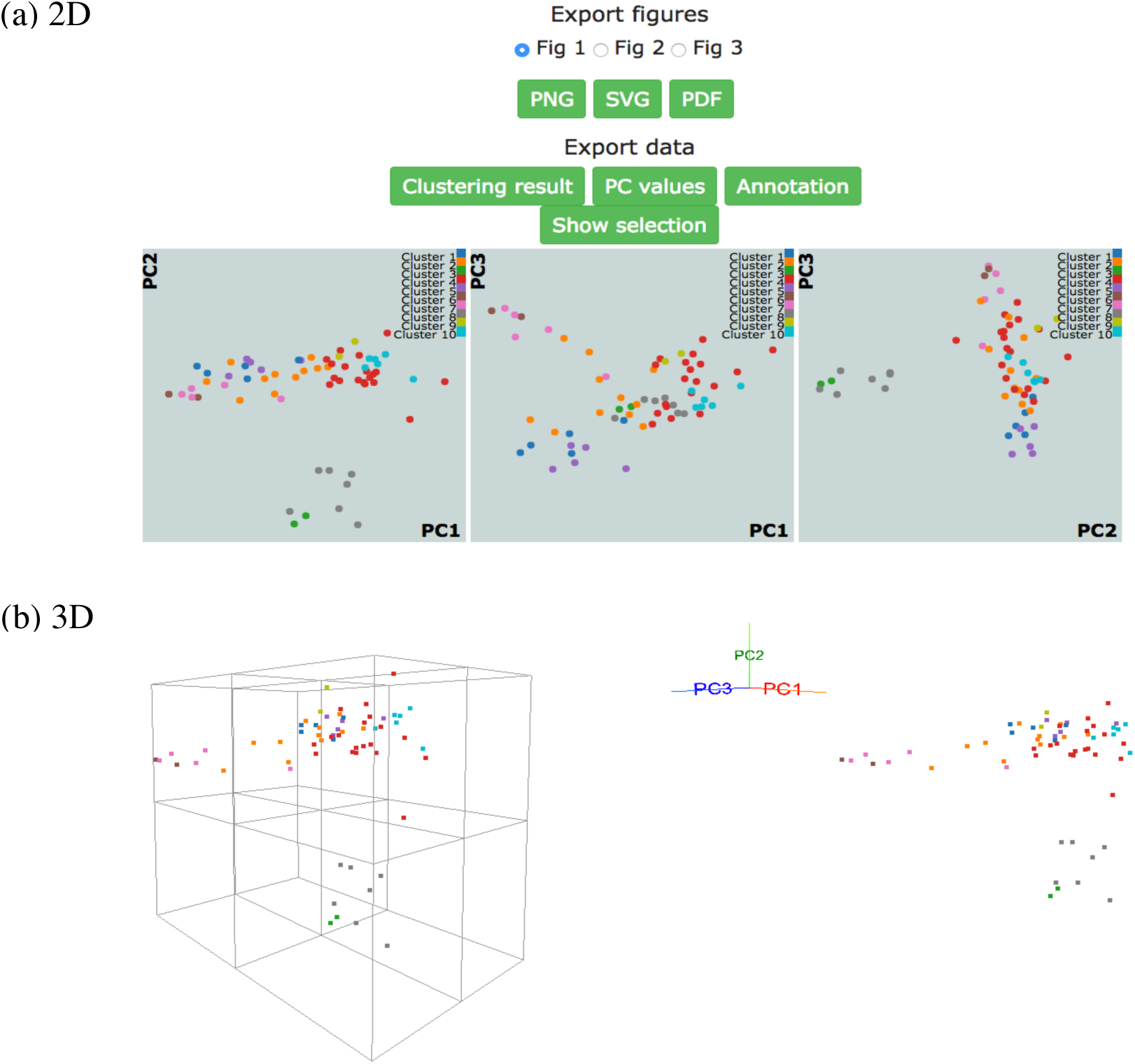
NCI-60 gene expression sample data clustering of samples using k-medoids algorithm, showing visualizations of first three principal components in (a) 2D and (b) 3D with perspective and orthogonal projection.

We also implement a dynamic 3D visualization for the first three principal components to enable deeper exploration of data structure by providing a perspective 3D view of data points. A real-time orthogonal projection from the current 3D viewpoint is also provided. Written in Javascript with libraries d3.js (Bostock et al., 2011), three.js (Cabello, 2010), the 3D Principal Component Viewer (Fig. 3b) allows zooming and rotating of the viewpoint. Graphical User Interface (GUI) is written using dat.GUI to toggle points or automate rotation of viewpoint. It should be noted that the user’s browser and machine capabilities may limit 2D and 3D visualizations. Our preliminary tests show that the visualizations work well for up to a few thousand data points on machines with typical hardware and modern browsers.

### Clustering tutorial and dynamic clustering

A detailed tutorial page on the website provides the user with a summary of advantages and disadvantages of each of the clustering algorithms. Interactive clustering from R Shiny package (Chang et al., 2015) is available for affinity propagation and Gaussian mixture model, allowing the user to add data points dynamically through the GUI and to observe changes in clustering behavior in real time (Fig. 4). The tutorial page further discusses pathological situations in which each algorithm may fail, with modified examples from the Scikit-learn Python package (Pedregosa et al., 2011).

**Fig. 4.**
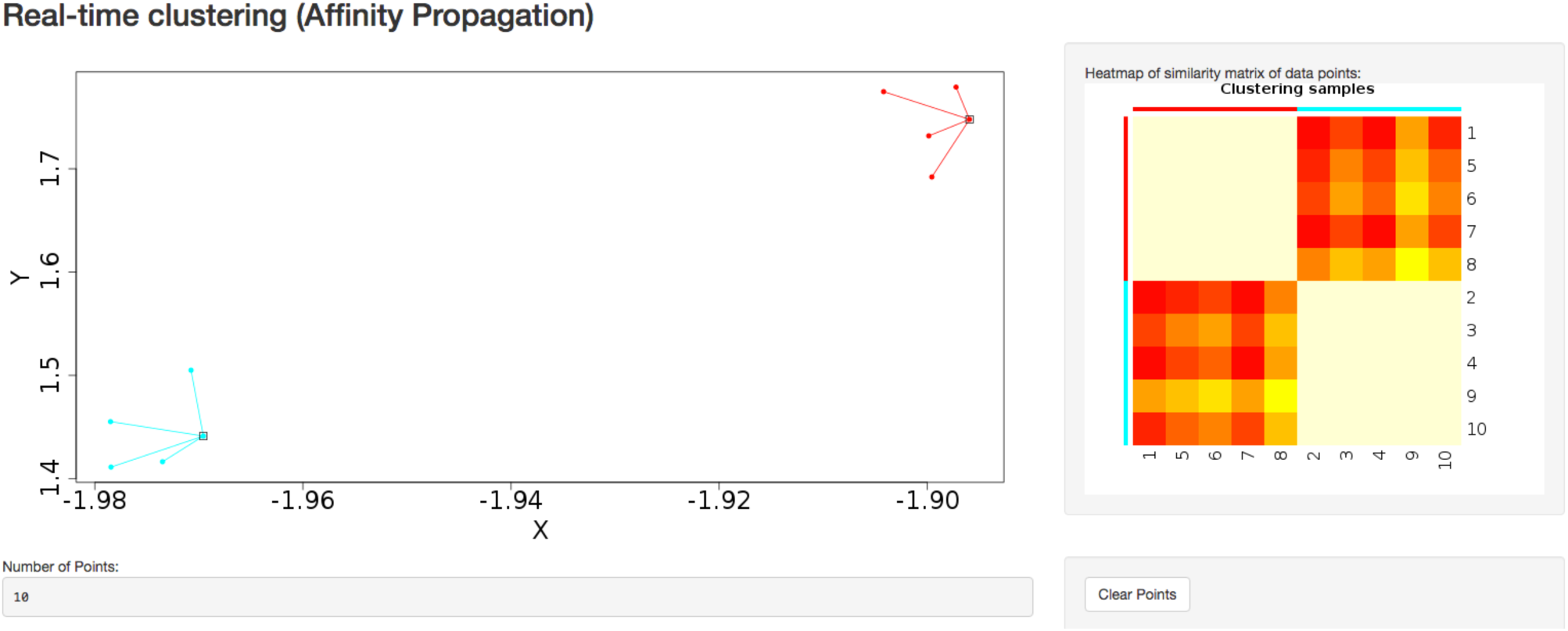
Dynamic clustering application in affinity propagation using R Shiny server displaying heatmap of similarity matrix of selected data points.

## 3 Results and Discussion

ClusterEnG offers a one-stop web service for efficient clustering of big data with the flexibility of choosing among many state-of-the-art clustering algorithms, which are not readily accessible to beginners. Interactive visualizations of clustering results in 2D and 3D enable users to comprehend their data effectively.

### Sample data

Fig. 3 shows snapshots of clustering results of NCI-60 sample data using k-medoids algorithm and 10 clusters. The samples are labeled using the same color scheme for both 2D and 3D visualizations. In Fig. 3a, k-medoids algorithm is able to separate closely related samples in terms of gene expression. For example, in the plot corresponding to PC1-PC2, the two breast tumor samples (green color) are identified in a cluster different from the nearby melanoma samples (gray color). In a similar way one can compare the clustering results across different algorithms and assess based on biological knowledge.

### Performance benchmarking results

We have benchmarked the performance of the codes for all available clustering algorithms. Fig. 5 shows the runtime of various clustering algorithms on ClusterEnG as a function of number of samples. The test data are randomly generated by using Gaussian distribution over the feature set for each sample. The number of features for each dataset is fixed at 10000, while the number of samples is varied from 100 to 5000. Fig. 5 also includes time taken to do PCA over the test data. The PCA step is common to all the algorithms. As we can observe from Fig. 5, DBSCAN performs best for all test data whereas affinity propagation and hierarchical clustering have maximum runtimes for larger and smaller sample sizes, respectively. In the above analysis, the parallel k-means and spectral clustering algorithms are run on a single core for comparison with serial codes.

**Fig. 5.**
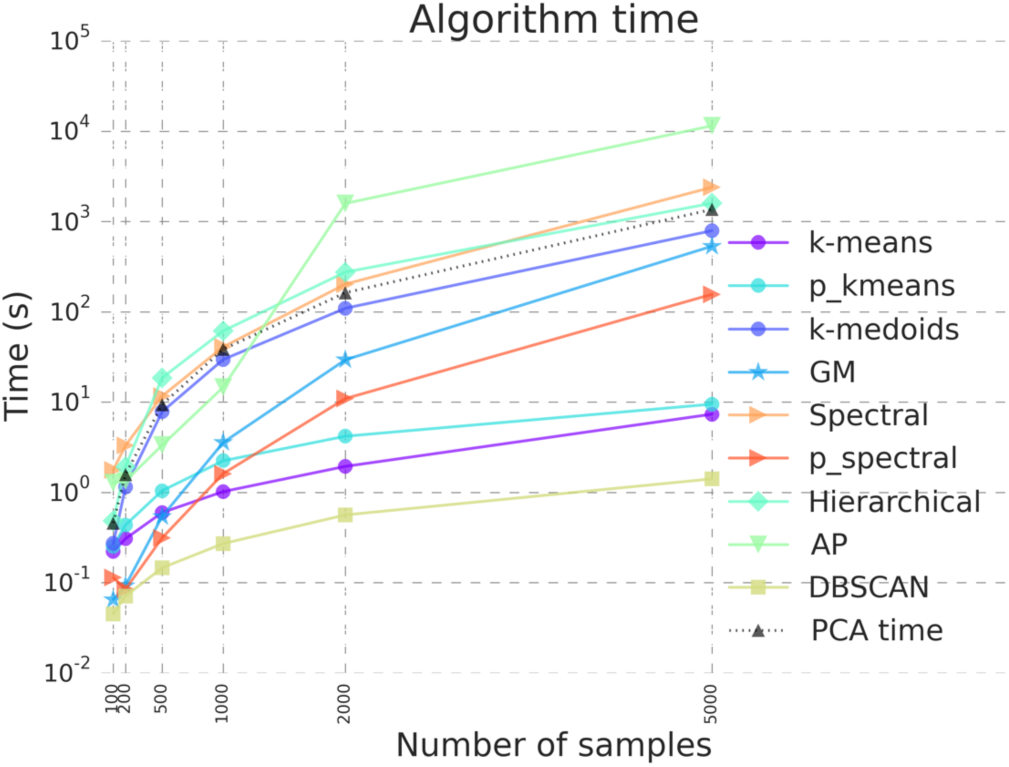
Benchmarking results illustrating algorithm run time for the clustering algorithms in ClusterEnG. “PCA time” data indicates time taken to compute principal components, a step common to all the algorithms for visualization.

We are currently developing and implementing parallel algorithms for affinity propagation and hierarchical clustering, and they will be included in the future releases of ClusterEnG. Furthermore, we plan to incorporate modules for exporting the clustering results directly to other available web servers for integrated analyses, including gene ontology and gene set enrichment analysis.

## Acknowledgements

The authors would like to thank the entire KnowEnG development team and the members of the Song group for valuable feedback and help.

